# Nitrogen-Responsive Extracellular Proteomics Reveals Evidence for a Novel Heterocyst-Specific Protein Secretion Pathway in *Anabaena*

**DOI:** 10.64898/2026.06.08.730779

**Authors:** Taufiq Nawaz, Ping He, Liping Gu, James Young, Ruanbao Zhou

**Affiliations:** Department of Biology and Microbiology, South Dakota State University, Brookings, 57007, SD, USA; BioNitrogen Economy Research Center, South Dakota, USA

**Author notes:** Corresponding authors: Liping Gu and Ruanbao Zhou. Guangdong Provincial Key Laboratory of Protein Function and Regulation in Agricultural Organisms, College of Life Sciences, South China Agricultural University, Guangzhou, Guangdong 510642, China (Ping He).

**Keywords:** Cyanobacteria, Heterocyst, Extracellular Proteome, Nitrogen Fixation, Non-Classical Secretion Pathways

## Abstract

Nitrogen availability is a major factor governing the physiology, ecology, and metabolism of cyanobacteria. Here, we performed a comparative extracellular proteomic analysis of *Anabaena* sp. PCC 7120 grown under nitrate-replete and diazotrophic conditions. Using LC–MS/MS, we identified 115 extracellular proteins in nitrate-grown cultures and 113 proteins under N_2_-fixing conditions. Remarkably, SignalP 6.0 predicted canonical signal peptides in only ∼22% of the identified proteins, suggesting that extracellular protein export in *Anabaena* predominantly occurs through non-classical secretion mechanisms, potentially involving extracellular vesicles or other unrecognized pathways.

Six highly abundant extracellular proteins (Alr2938, Alr4550, Alr2328, All4121, Alr0528, and Alr0529) were detected under both nitrogen regimes. In contrast, All4337, Alr0608, and All3093 were preferentially enriched under nitrate-replete conditions, whereas Alr0267 and Alr1050 emerged among the most abundant extracellular proteins during diazotrophic growth. Notably, an Alr0267-GFP fusion protein was detected exclusively in heterocysts, the specialized N_2_-fixing cells of *Anabaena*, with GFP fluorescence concentrated at the cell periphery. The extracellular localization of Alr0267 is particularly intriguing because heterocysts are surrounded by specialized polysaccharide and glycolipid envelope layers that establish the microoxic environment required for nitrogenase activity. The apparent export of Alr0267 across these barriers provides evidence for a previously unrecognized heterocyst-associated protein secretion pathway.

Together, these findings reveal a nitrogen-responsive extracellular proteome and provide the first evidence for heterocyst-specific extracellular protein secretion. This work advances our understanding of heterocyst biology and protein trafficking while laying a foundation for engineering *Anabaena* as a sustainable photosynthetic platform for secreting high-value proteins using sunlight, CO_2_, N_2_, and mineralized water.

## 1. Introduction

*Anabaena* sp. PCC 7120 is a filamentous diazotrophic cyanobacterium that has served as a premier model organism for investigating nitrogen fixation, cellular differentiation, and multicellular organization in prokaryotes. In response to combined nitrogen deprivation, approximately 5–10% of vegetative cells differentiate into heterocysts, specialized terminally differentiated cells dedicated to biological nitrogen fixation (1, 2). This developmental transition is essential because the nitrogenase enzyme complex responsible for atmospheric N_2_ reduction is highly sensitive to oxygen. To maintain the microoxic conditions required for nitrogenase activity, heterocysts develop a specialized envelope consisting of an inner glycolipid layer and an outer polysaccharide layer that effectively restrict oxygen diffusion (3). Owing to its fully sequenced genome, genetic tractability, and well-characterized developmental program, *Anabaena* sp. PCC 7120 has become an important model system for elucidating the molecular mechanisms governing nitrogen fixation and cellular differentiation in cyanobacteria.

In addition to their intracellular metabolic activities, cyanobacteria actively interact with their environment through the secretion of extracellular proteins. These proteins participate in diverse biological processes, including nutrient acquisition, cell signaling, biofilm formation, extracellular matrix remodeling, and environmental adaptation (4-7). The cyanobacterial cell envelope is a complex structure composed of peptidoglycan, outer membrane proteins, surface-associated pigments, and extracellular polysaccharides, all of which contribute to cellular interactions with the surrounding environment (8). Protein secretion in cyanobacteria has traditionally been associated with the Sec and Tat pathways. For example, Sergeyenko and Los (2000) identified eight secreted proteins in *Synechocystis* sp. PCC 6803, five of which contained putative signal peptides characteristic of canonical secretion systems (7). However, accumulating evidence suggests that extracellular protein export in cyanobacteria may involve additional mechanisms that remain poorly understood.

Several studies have examined the extracellular proteome of *Anabaena* sp. PCC 7120 under nitrate-containing growth conditions. Comparative exoproteomic analyses revealed altered secretome profiles in mutants defective in the TolC-like outer membrane channel HgdD (9), the zinc-responsive regulator Zur (10), and the global regulator FurC (11), highlighting the importance of extracellular proteins in nutrient acquisition and environmental adaptation. More recently, a comprehensive exoproteomic survey identified 139 extracellular proteins in *Anabaena* cultures grown with nitrate, ammonium, or under nitrogen-fixing conditions (12). Despite these advances, the impact of nitrogen availability on extracellular protein composition and secretion mechanisms remains incompletely understood. In particular, little is known about whether heterocysts contribute directly to extracellular protein secretion and how proteins could be exported across the specialized glycolipid and polysaccharide envelope layers that surround these heterocysts, the nitrogen-fixing cells.

Given the central role of nitrogen metabolism in *Anabaena* physiology, characterizing the extracellular proteome under contrasting nitrogen regimes may provide new insights into cyanobacterial adaptation and cell differentiation. In this study, we performed a comparative extracellular proteomic analysis of *Anabaena* sp. PCC 7120 grown under nitrate-replete (AA/8N) and diazotrophic (AA/8) conditions. We identified 115 and 113 extracellular proteins, respectively, and found that the majority lacked canonical secretion signal peptides, suggesting an important role for non-classical secretion pathways. Most notably, we identified Alr0267 as a highly abundant extracellular protein under diazotrophic conditions and demonstrated its specific localization to heterocysts. These findings provide evidence for heterocyst-associated extracellular protein secretion and raise the possibility of a previously unrecognized protein export pathway capable of transporting proteins across the specialized heterocyst envelope. Beyond advancing our understanding of heterocyst biology and cyanobacterial protein trafficking, this work provides a foundation for engineering *Anabaena* as a sustainable photosynthetic platform for the extracellular production of recombinant proteins and other high value bioproducts using sunlight, CO_2_, N_2_, and mineralized water.

## 2. Materials and methods

### 2.1. Bacterial Strain and Culture Conditions

*Anabaena sp*. PCC 7120 cultures were maintained at 30 ± 2 °C in 1/8 strength modified Allen-Arnon medium [AA/8N: with combined nitrogen (10 mM NaNO_3_) or AA/8: nitrogen-free] (13) under continuous illumination at an intensity of 70 µmol photons m^-2^ s^-1^ using cool-white fluorescent lamps (14, 15).Cultures were grown in Erlenmeyer flasks with constant agitation at 110 rpm to ensure homogeneous aeration and mixing. All experiments were performed under axenic conditions (16, 17).

### 2.2. Extracellular Protein Isolation

Wild-type *Anabaena sp*. PCC 7120 was initially cultured in 30 mL volumes of either AA/8N or AA/8 medium until reaching an optical density of 0.3–0.4 at 720 nm. Cultures were centrifuged at 8,000 × g for 8 min at 20 °C to pellet the cells. The pellets were washed three times with sterile nitrogen-free AA/8 medium to remove any residual extracellular proteins. After washing, the cells were resuspended in 1 mL of AA/8 medium and inoculated into 1 L of fresh AA/8N or AA/8 medium in glass flasks. Cultures were incubated under the same light and temperature conditions for 10 days to allow extracellular protein accumulation (18). Following incubation, cells were separated from the culture supernatant by centrifugation at 10,000 × g for 20 min at 20 °C. The resulting supernatants were immediately filtered through 0.22 μm pore size nitrocellulose membranes to remove any remaining cell debris. Extracellular proteins were precipitated by adding solid ammonium sulfate to a final concentration of 80% (w/v) and incubated overnight at 4 °C with gentle stirring (19). Precipitated proteins were collected by centrifugation at 10,000 × g for 40 min at 4 °C. Protein pellets were resuspended in 3–4 mL of 10 mM phosphate buffer (pH 7.4) and dialyzed against deionized water for 12–24 h at 4 °C using dialysis tubing with a 10 kDa molecular weight cut-off (20). The dialysis water was replaced 4–5 times to ensure complete removal of salts. The dialyzed protein solution was then concentrated by vacuum freeze-drying and stored at –20 °C until further analysis.

### 2.3. SDS-PAGE and In-Gel Digestion for Mass Spectrometry

Lyophilized extracellular protein powders were resuspended in 120 μL of Laemmli SDS-PAGE loading buffer, boiled at 95 °C for 8 min to denature proteins, and briefly centrifuged at 10,000 × g for 1 min to remove insoluble particulates. The supernatants were directly loaded onto 12% SDS-PAGE gels and electrophoresed at room temperature (21). Following electrophoresis, proteins were visualized by staining with 0.1% Coomassie Brilliant Blue R-250 in 45% methanol and 10% acetic acid for 1 h, followed by destaining overnight in a solution containing 20% methanol and 10% acetic acid (22). For specific band protein identification under AA/8N and AA/8 condition, equal amounts of protein (∼60 μg per lane) were run into a 12% polyacrylamide SDS-PAGE gel (Bio-Rad The PROTEAN II xi cell, 1.5 mm thick) at 200 V for approximately 2 hour and specific bands were excised for tryptic digestion and LC-MS/MS analyses (Figure 1). As for total protein identification under AA/8N and AA/8 condition, equal amounts of protein were run into a 12% polyacrylamide SDS-PAGE gel (Bio-Rad Mini-PROTEAN, 1.0 mm thick) at 200 V for ∼15 min until the protein entered the resolving gel (the front dye migrated into resolving gel for approximately 1 cm). The gel was stained with Coomassie Brilliant Blue R-250, and each lane was excised as a single band (Figure 2).

**Figure 1.**
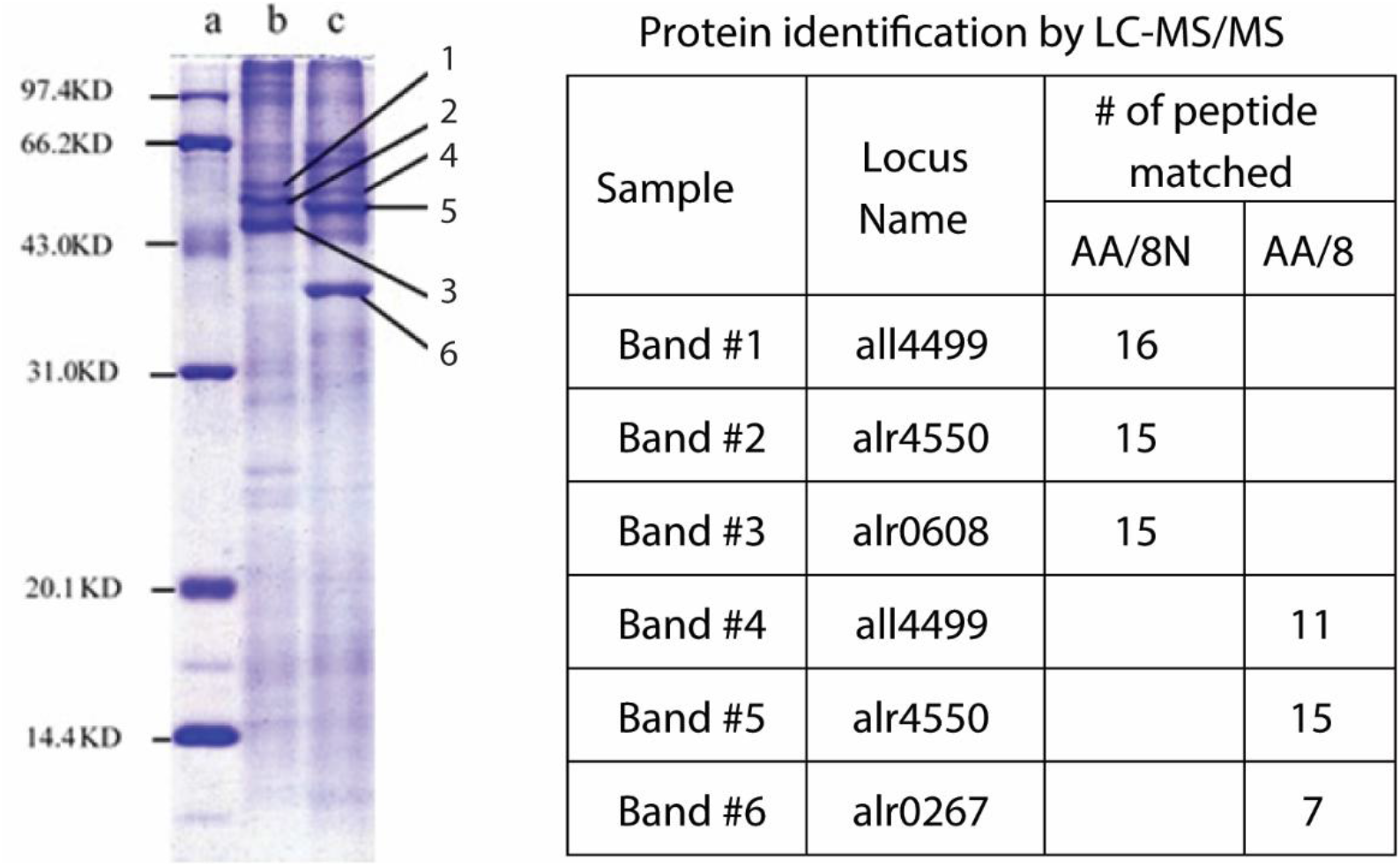
Extracellular proteins identified from AA/8N or AA/8 culturing medium of *Anabaena sp*. Strain PCC 7120. Exoproteome patterns of *Anabaena sp*. PCC 7120 cultivated under AA/8 and AA/8N growth conditions. (**a**) Protein Ladder; (**b**) Extracellular proteins extracted from AA/8N culturing medium after centrifugation of removing the cell pellet and [NH_4_]_2_SO_4_ precipitation to concentrate and fractionate proteins from extracellular matrix; and (**c**) Extracellular extracted from AA/8 culturing medium. Bands 1-6 were selected for in-gel tryptic digestion and protein identification by mass spectrometry. The number of peptides matched to the identified proteins through LC-MS/MS was summarized. (for detail information, see Supplementary Information S1).

**Figure 2.**
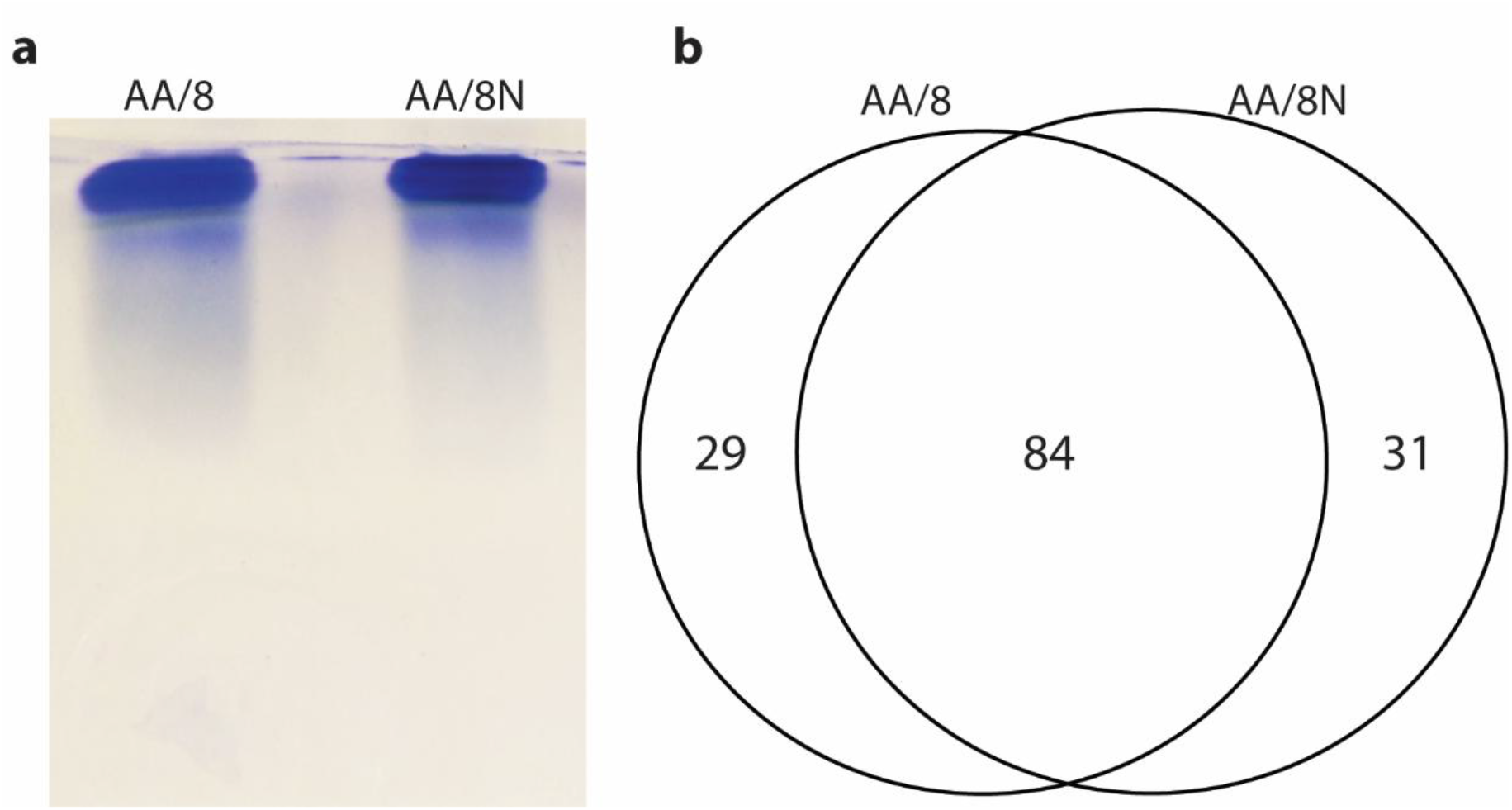
Total extracellular proteins identified from AA/8N or AA/8 culturing medium of *Anabaena sp*. Strain PCC 7120. (**a**) Total extracted extracellular proteins concentrated by 12% SDS-PAGE (run at 200 mV for approximately 15 min until all the proteins just entered the resolving gel) and (**b**) Venn diagram showed a total of 113 and 115 proteins were identified from AA/8 and AA/8N media by LC-MS/MS, respectively.

### 2.4. In-gel tryptic digestion and protein identification by LC-MS/MS

The protein gel bands (Figure 1 and Figure 2) were excised and in-gel tryptic digestion was performed according to Shevchenko (Shevchenko et al., 1996) with the following modifications. Briefly, gel slices were dehydrated with acetonitrile (ACN) for approximately 5 min incubation and repeated this process until they appear to shrink in size and show a chalk white color. The time required and number of washes vary with gel size and composition. The chalk white color gel was then incubated with 100 mM ammonium bicarbonate (NH_4_HCO_3_) containing 10 mM dithiothreitol (DTT, pH ≈ 8.0) for 45 min at 56°C, dehydrated again and incubated with 100 mM NH_4_HCO_3_ containing 50 mM iodoacetamide for 20 min in the dark, and then washed with 100 mM NH_4_HCO_3_ and dehydrated again. Approximately 50 μL trypsin solution (0.01 μg/μL sequencing grade modified trypsin (Promega #V5111) in 50 mM NH_4_HCO_3_) was added to each gel slice so that the gel was completely submerged and then incubated at 37°C for overnight. The tryptic peptides were extracted with 60% ACN/1% TCA from the gel by water bath sonication (Aquasonic 150T sonicating water bath which puts out 135W. Sonication is done 2 × 20s) and concentrated in a SpeedVac to 5 μL.

For specific band protein identification (specific bands labeled 1 to 6 in Figure 1), 5 μL of the extracted peptide suspension was injected (to the sample loop which is then backflushed using solvent A directly to the column) by EASYnLC and the peptides separated through an Acclaim PepMap RSLC column (0.075 mm x 150 mm C18, Thermo Scientific) using a nanoAcquity UPLC (Waters) (Buffer A = 99.9% Water/0.1% Formic Acid, Buffer B = 99.9% Acetonitrile/0.1% Formic Acid). Peptides were eluted with a linear gradient from 5 to 30% B developed over 35 min. For total protein identification (condense protein bands shown in Figure 2), peptides were eluted with a linear gradient from 5 to 30% B developed over 120 min. The eluted peptides were sprayed into a Q Exactive hybrid quadrupole-Orbitrap mass spectrometer using a Nanospray Flex™ Ion Sources (Thermo Scientific). Survey scans were taken in the Orbi trap (35,000 resolution determined at *m/z* 200) and the top ten ions in each survey scan were then subjected to automatic higher energy collision induced dissociation (HCD) with fragment spectra acquired at 17,500 resolutions (by convention this is a dimensionless measurement).

For protein identification, the resulting MS/MS spectra were converted to peak lists using Mascot Distiller (www.matrixscience.com) and searched against a protein sequence database containing *Anabaena* (*Nostoc*) *sp*. PCC 7120 and common laboratory contaminants downloaded from www.thegpm.org. All searches were performed using the Mascot searching algorithm, v 2.4. The Mascot output was then analyzed using Scaffold (www.proteomesoftware.com) to probabilistically validate protein identifications filtered with 95% minimum protein probability, 2 minimum identified peptides, and 95% minimum peptide probability. The putative signal peptide was analyzed using SignalP 6.0 (https://services.healthtech.dtu.dk/services/SignalP-6.0). Functional categorization of the exoproteome according to the COG classification provided by EGGNOG mapper V2 available at https://github.com/eggnogdb/eggnog-mapper. Prediction of subcellular localization using PSORTb software version 3.0.3 available at https://www.psort.org/psortb/ and DeepLocPro - 1.0 available at https://services.healthtech.dtu.dk/services/DeepLoc-1.0/

### 2.5. Plasmid construction and subcellular localization of Alr0267-GFP fusion protein

A 2075 bp fragment (alr0267 ORF along with its native promoter region) was PCR-amplified by Phusion™ High-Fidelity DNA Polymerase (NEB) from genomic DNA of *Anabaena* 7120 using the primers ZR1213 (5’-t<u>ctcgagg</u>ATCCCTGTACCACTACCAATCTTCG-3’) and ZR1230 (5’-t<u>cccggg</u>AAGCTGTGTACCACTCTGGAGTC-3’) and cloned into pTOPO2.1 vector to produce pZR2071; Then the 2091 bp fragment from pZR2071 was cut out by BamHI + XmaI and ligated into BamHI + XmaI digested pSMC232-GFP vector to produce pZR2099 (illustrated in Figure S1). The PCR verified pZR2099 was transformed into *Anabaena sp*. PCC 7120 to create *Anabaena* strain A2099 bearing Alr0267-GFP through a conjugation method (23). Alr0267-GFP localization was monitored using Olympus BX53 Fluorescence Microscope equipped with GFP-4050A filter (BrightLine® EX466/40 nm, EM525/50 nm, DM495).

## 3. Results and discussions

### 3.1 *Anabaena sp*. PCC 7120 Exoprotein Identification

To explore the exproteomes in the *Anabaena sp*. PCC 7120, we first grew *Anabaena sp*. PCC 7120 in nitrogen-free and nitrate-supplied medium, collected the supernatant and precipitated the extracellular proteins with ammonium sulfate to concentrate and fractionate proteins from the extracellular matrix, then performed SDS-PAGE separation, band excision, in-gel tryptic digestion, and LC-MS/MS analysis. As shown in Figure 1, three protein bands from each line (highlighted by arrowheads) were subjected for further LC-MS/MS protein identification (See Supplementary Information S1: Mascot Search Results for the specific protein bands in Figure 1). All4499, Alr4550 were detected in nitrogen-fixing conditions (AA/8) or in medium supplemented with nitrate AA/8N, consistent with the previous report that both All4499 and Alr4550 were detected in BG-11_0_ (11) and BG-11 growth media (9, 11). The third protein band in medium supplemented with nitrate AA/8N was identified as nitrate transport nitrate-binding protein Alr0608, in agreement with this protein function as nitrate-binding and transporting. Alr0267 (heterocyst-specific attachment factor or HesF) was detected in nitrogen-fixing conditions (AA/8), consistent with its up-regulation in response to nitrogen deprivation (24, 25). Taken together, the above approaches allow identifying the presence or absence of the proteins extracted from the culturing media with or without nitrate.

Since the above approach that does not allow performing direct comparisons of exoproteome differential compositions between culturing conditions, a modified method was adapted by running the protein samples just into resolving gel of the SDS-PAGE for purification, excising the bands for tryptic digestion, and then performing LC-MS/MS proteome analysis as illustrated in Figure 2. The Venn diagram represents the unique and shared proteins (113 and 115 proteins were identified in AA/8 and AA/8N culturing conditions, respectively (See Supplementary Information S2: Protein and Peptide Reports for the condensed protein bands in Figure 2 and Table S1: Overview of the identified proteins).

When compared the set of proteins identified here to previously reported exoproteomes of *Anabaena* under similar growth conditions, we found that among 113 proteins identified in AA/8 in this study, 44 was previously identified in nitrogen-fixing conditions (BG-11_0_) by Oliveira and colleagues (11). Similarly, among 115 proteins identified in AA/8N in this study, 36.5% of the proteins was identified by Oliveira and colleagues (11), 22.6% of the proteins from wild-type *Anabaena sp*. PCC 7120 was found in the Hahn exoproteome (26) and 78.3%) of the proteins in the Sarasa-Buisan exoproteome under in medium supplemented with nitrate (BG-11) (9, 10) (Table S2).

Among 144 proteins identified, 29 proteins were only found in nitrogen-fixing conditions (AA/8) (Table 1) and 31 found in medium supplemented with nitrate AA/8N (Table 2). The relative abundance of the proteins was assessed using the Scaffold software to calculate the quantitative value number by normalizing spectral counts across samples.

**Table 1.** Unique Proteins identified in culturing medium without combined nitrogen (AA/8)

| **Locus Name** | **Protein Description** | **Quantitative Value** | **# of Assigned Spectra** | **# of Unique Spectra** | **# of Unique Peptides** |
| --- | --- | --- | --- | --- | --- |
| alr0267 | hypothetical protein | 47.036 | 53 | 11 | 9 |
| alr1050 | pgi gene product | 39.049 | 44 | 19 | 17 |
| alr7163 | transposase | 11.537 | 13 | 4 | 2 |
| alr4777 | hypothetical protein | 7.9873 | 9 | 5 | 4 |
| all0875 | hypothetical protein | 6.2124 | 7 | 3 | 3 |
| alr1077 | hypothetical protein | 6.2124 | 7 | 4 | 4 |
| alr3932 | hypothetical protein | 6.2124 | 7 | 3 | 3 |
| alr4853 | aspartate aminotransferase | 5.3249 | 6 | 4 | 4 |
| all1430 | Heterocyst Ferredoxin | 5.3249 | 6 | 3 | 2 |
| all4388 | hypothetical protein | 5.3249 | 6 | 4 | 4 |
| all2105 | acpD gene product | 4.4374 | 5 | 4 | 4 |
| all2533 | prolyl endopeptidase | 4.4374 | 5 | 3 | 3 |
| all4749 | hypothetical protein | 4.4374 | 5 | 3 | 3 |
| alr0996 | protease | 4.4374 | 5 | 2 | 2 |
| all0955 | hypothetical protein | 3.5499 | 4 | 3 | 3 |
| alr1310 | hypothetical protein | 3.5499 | 4 | 2 | 2 |
| all0458 | hypothetical protein | 3.5499 | 4 | 3 | 2 |
| all2303 | dihydroorotase | 2.6624 | 3 | 2 | 2 |
| alr3608 | hypothetical protein | 2.6624 | 3 | 2 | 2 |
| all1380 | hypothetical protein | 2.6624 | 3 | 2 | 2 |
| alr2405 | Flavodoxin | 2.6624 | 3 | 2 | 2 |
| alr3339 | dihydroorotase | 2.6624 | 3 | 2 | 2 |
| all4089 | hypothetical protein | 2.6624 | 3 | 2 | 2 |
| asl4547 | hypothetical protein | 2.6624 | 3 | 2 | 2 |
| alr1594 | DNA-directed RNA polymerase beta subunit | 1.775 | 2 | 2 | 2 |
| alr1622 | hypothetical protein | 1.775 | 2 | 2 | 2 |
| alr2887 | hypothetical protein | 1.775 | 2 | 2 | 2 |
| alr3683 | ATP-dependent Clp protease proteolytic subunit 3 | 1.775 | 2 | 2 | 2 |
| all4555 | hypothetical protein | 1.775 | 2 | 2 | 2 |

**Table 2.**
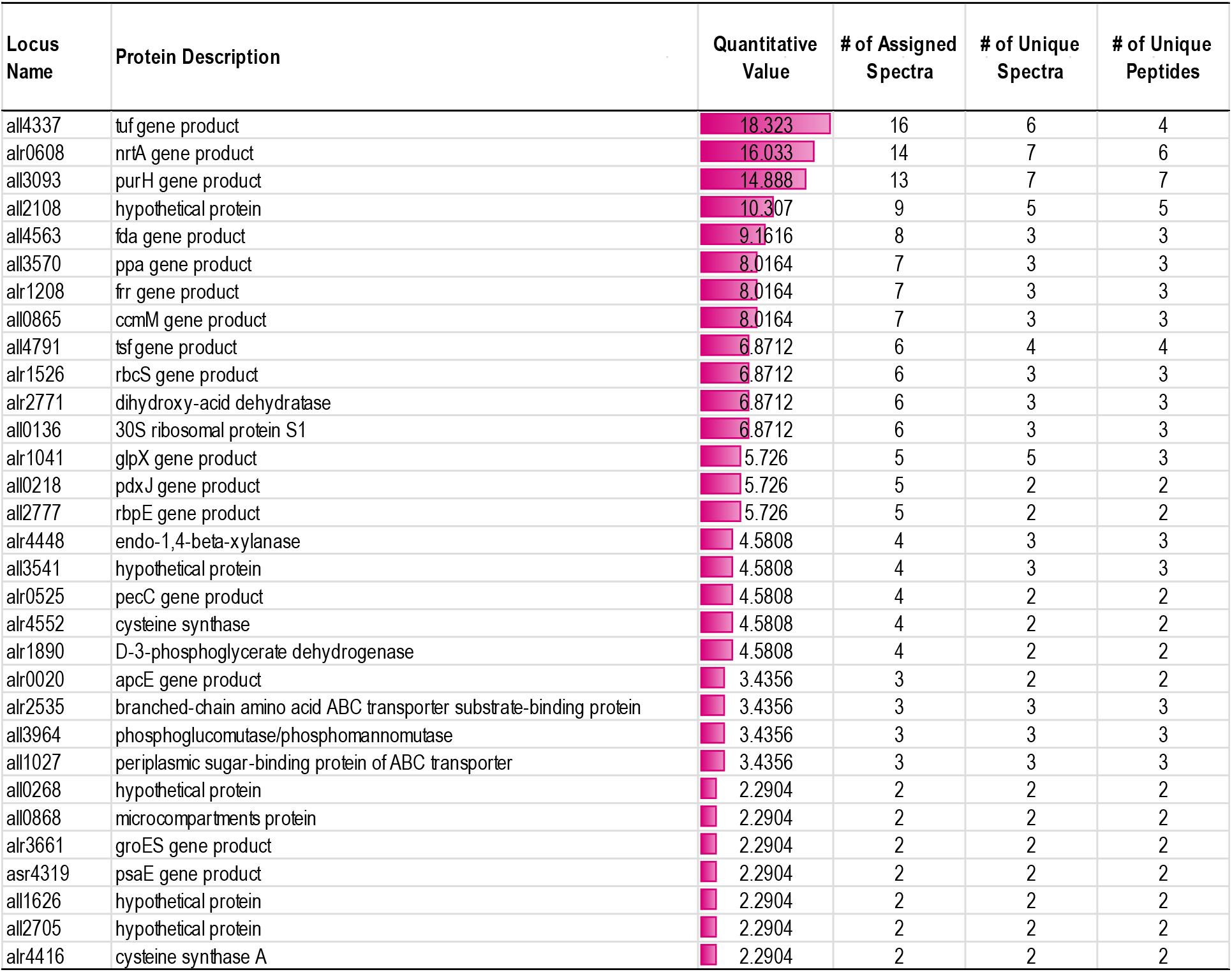
Unique Proteins identified in culturing medium with combined nitrogen (AA/8N)

### 3.2. Unique Proteins identified in culture medium without combined nitrogen

There are 29 unique proteins identified in *Anabaena sp*. PCC 7120 grown diazotrophically (Table 1), which were annotated to participate in cellular processes and signaling (Alr0996 and Alr3683), amino acid transport and metabolism (Alr4853 and All2533), carbohydrate transport and metabolism (Alr1050, All0875, and Alr1310), energy production and conversion (All1430 and Alr2405), inorganic ion transport and metabolism (All0458, Alr3608, and All4555), nucleotide transport and metabolism (All2303 and Alr3339), and secondary metabolites biosynthesis, transport, and catabolism (Alr0267, Alr1077, and All0955), and cell wall/membrane/envelope biogenesis (All4388 and Alr2887). Consistent with our observation, Oliveira et. al. reported that All4388 was only detected in nitrogen-free medium (11). All4388 is a Fox gene that encodes a putative polysaccharide transporter and required for heterocyst envelope polysaccharide deposition (27).

Alr0267 was previously reported to be found in the extracellular milieu of *Anabaena sp*. PCC 7120 grown in BG-11_0_ (11) and BG-11 (9, 11, 26). Alr0267, also known as HesF (heterocyst-specific attachment factor), was further characterized using GFP as a reporter and showed that the alr0267 promoter was specifically activated in heterocysts (25), in an agreement with the observation that the alr0267 transcripts was up-regulated in response to nitrogen deprivation (24, 25). To determine the subcellular localization of HesF, we constructed pZR2099 plasmid containing P_alr0267_-Alr0267-GFP for Alr0267-GFP expression driven by alr0267 native promoter (Figure 3a and Figure S1). As shown in Figure 3b, Alr0267-GFP fusion protein was only detected in heterocysts of *Anabaena sp*. PCC 7120 grown diazotrophically in AA/8 medium and the fluorescence signals were revealed to be enriched at the heterocyst periphery. Taken together, these data suggest that HesF was transcripted and translated solely within heterocysts and released into the extracellular milieu of *Anabaena sp*. PCC 7120 grown diazotrophically.

**Figure 3.**
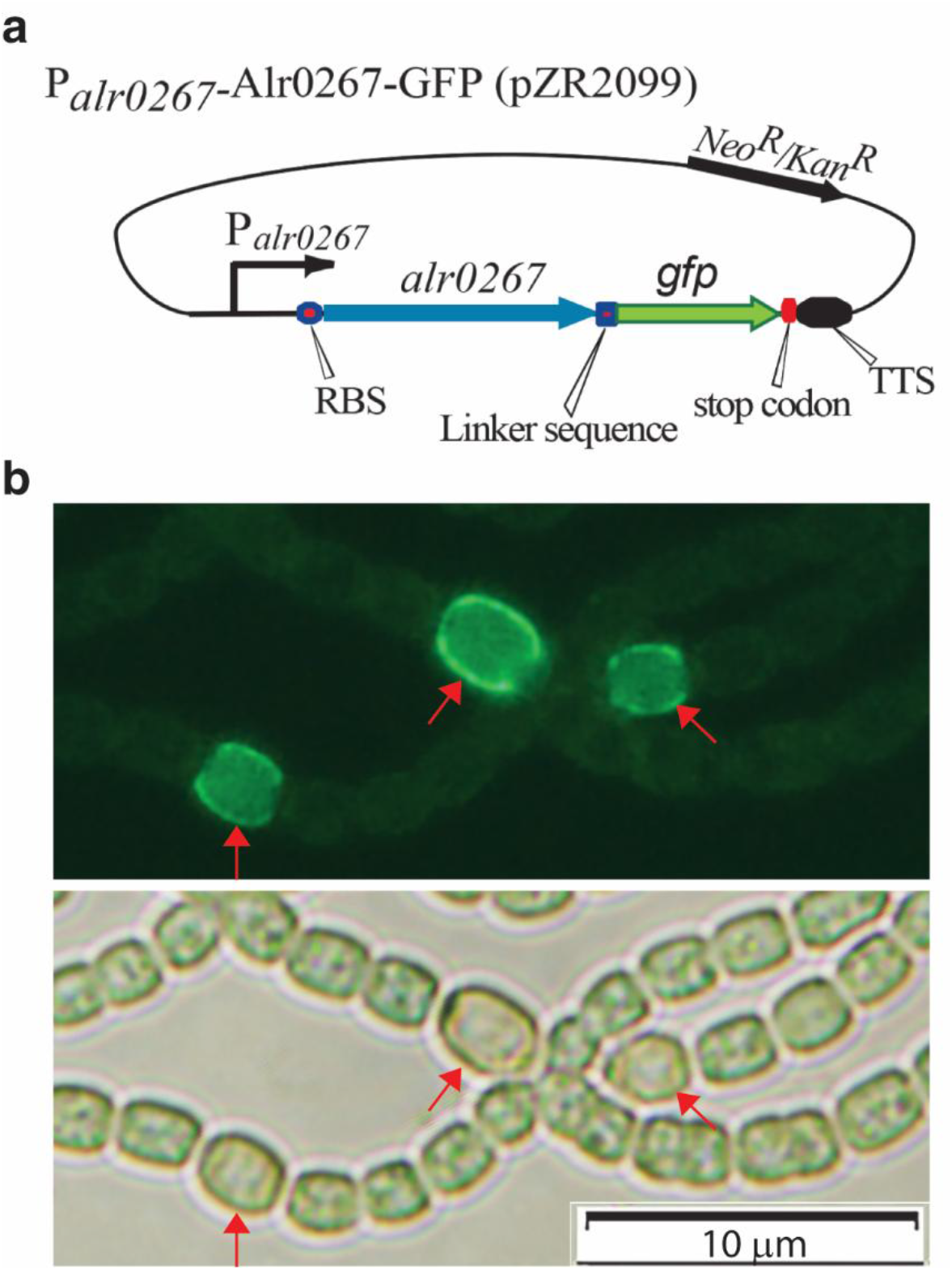
Subcellular localization of Alr0267-GFP in *Anabaena sp*. PCC 7120 grown diazotrophically in AA/8 medium. **a.** Schematic representation of the construction of pZR2099 for expressing the Alr0267-GFP fusion protein driven by the alr0267 native promoter. **b**. The Alr0267-GFP fusion protein was only detected in heterocysts of *Anabaena sp*. PCC 7120 and the fluorescence signals were enriched at the periphery (*top panel*: GFP filter; *bottom panel*: Brightfield).

To our knowledge, this study provides the first evidence for heterocyst-specific extracellular secretion of Alr0267 in *Anabaena sp*. PCC 7120. Among the proteins identified in the extracellular proteome, Alr0267 was particularly noteworthy because it was highly abundant under diazotrophic conditions and its expression was restricted to heterocysts, the specialized nitrogen-fixing cells responsible for biological N_2_ fixation. Fluorescence microscopy further revealed that Alr0267-GFP was concentrated at the periphery of heterocysts, consistent with a role associated with the cell envelope or extracellular environment.

The apparent extracellular localization of Alr0267 is especially intriguing in light of the unique architecture of heterocysts. Unlike vegetative cells, heterocysts develop specialized polysaccharide and glycolipid envelope layers that form an effective barrier to oxygen diffusion and create the microoxic conditions required for nitrogenase activity. While these envelope layers have been extensively studied for their role in protecting nitrogenase, little is known about how proteins are transported across these specialized structures. The detection of Alr0267 in the extracellular proteome therefore raises a fundamental biological question: how can proteins produced within heterocysts traverse multiple envelope barriers and be released into the extracellular milieu?

Our findings suggest the existence of a previously unrecognized protein export mechanism associated with heterocyst differentiation and nitrogen fixation. This hypothesis is further supported by the observation that only a small fraction of the extracellular proteins identified in this study contained canonical Sec- or Tat-dependent signal peptides. The predominance of proteins lacking recognizable secretion signals suggests that non-classical secretion pathways may contribute substantially to extracellular protein trafficking in *Anabaena*. Potential mechanisms include extracellular vesicle-mediated transport, specialized envelope-associated secretion systems, or other unconventional export pathways that remain to be characterized.

The discovery of extracellular Alr0267 associated with heterocysts opens new avenues for investigating protein trafficking in differentiated cyanobacterial cells. Elucidating the molecular machinery responsible for Alr0267 export may uncover a previously unknown secretion pathway operating in heterocysts and provide new insights into the coordination of cellular differentiation, nitrogen fixation, and extracellular communication. Beyond its fundamental biological significance, such a secretion system could offer unique opportunities for engineering *Anabaena* as a solar-powered, sustainable microbial platform for the extracellular production of recombinant proteins using atmospheric dinitrogen gas (N_2_) as a sole nitrogen source.

The second most abundant exoprotein Alr1050 was solely identified in the culture of *Anabaena* grown diazotrophically with AA/8 medium (Table 1). This Alr1050 encoding glucose-6-phosphate isomerase involved in carbohydrate metabolism, which was previously found in *Anabaena sp*. PCC 7120 grown in both combined nitrogen-containing media and diazotrophic condition (11, 26), this inconsistence could possibly due to the differences of the growth conditions, exoprotein extraction methods, and the limit of detection of the LC-MS/MS instrumentations. Moreover, computational prediction analyses indicated its cytoplasmic localization without putative signal peptide (Table S3). Further investigation is needed to explore its secretion pathway, potential extracellular or structural roles, and its function in regulating carbon and nitrogen balance within *Anabaena*’s cellular metabolism.

### 3.3. Unique Proteins identified in culture medium with combined nitrogen (AA/8N)

There are 31 unique proteins identified in *Anabaena sp*. PCC 7120 grown diazotrophically (Table 2), which were annotated to participate in cellular processes and signaling (All1626, Alr3661, and, Asr4319), amino acid transport and metabolism (Alr1890, Alr4552, Alr2535, Alr4416, and Alr2771), carbohydrate transport and metabolism (All4563, Alr1041, Alr4448, All1027, All3964, and Alr0020), energy production and conversion ; secondary metabolites biosynthesis, transport and catabolism (All0865, All3570, Alr1526, and All0868), inorganic ion transport and metabolism (Alr0608), nucleotide transport and metabolism (All3093), and secondary metabolites biosynthesis, transport, and catabolism (Alr0267, Alr1077, and All0955). Furthermore, there were two categories that uniquely present in this group were proteins involved in translation, ribosomal structure, and biogenesis (All4337, Alr1208, All0136, and All4791) and coenzyme transport and metabolism (All0218 and Alr0525).

The most abundant exoproteins uniquely detected in *Anabaena sp*. PCC 7120 grown in the nitrate-containing medium were All4337, Alr0608, and All3093.

All4337 encodes the essential Translation Elongation Factor EF-Tu, a core protein responsible for delivering aminoacyl-tRNAs to the ribosome during protein biosynthesis. All4337, along with Alr1208, All0136, and All4791, may play an important role in response to RNA binding, protein biogenesis, and extracellular stress.

In contrast to the previous reports that Alr0608 was found as an exoprotein under the growth conditions with or without combined nitrogen (Table S2) (9-11, 26), this study only detected Alr0608 in *Anabaena sp*. PCC 7120 grown in AA/8N. Alr0608 encodes NrtA, which is the periplasmic nitrate/nitrite-binding protein component of the ATP-binding cassette (ABC) nitrate transport system (28, 29). NrtA gene is required for growth on nitrate and heterocyst development (30). Since nrtABCD are in an operon and function synergistically, it will be interested to further investigate NrtA as an exoprotein and its function in the extracellular milieu of *Anabaena sp*. PCC 7120.

The third most abundant exoprotein uniquely found in *Anabaena sp*. PCC 7120 grown in AA/8N medium was All3093, encoding bifunctional purine biosynthesis protein PurH, which was also detected by Sarasa-Buisan et. al. in *Anabaena sp*. PCC 7120 grown in BG-11 medium (9), suggesting its potential role in nucleotide transport and metabolism.

### 3.4. Differentially abundant exoproteins identified in culture media with and without combined nitrogen

There are 24 most abundant shared proteins (Quantitative Value >15) identified in *Anabaena sp*. PCC 7120 (Table 3), which were annotated to participate amino acid transport and metabolism (Alr2328, and All1951), carbohydrate transport and metabolism (All3538), coenzyme transport and metabolism (All2315), energy production and conversion (Alr2938, Alr0528, Alr0529, All4968, All5039, Alr0022, Alr0021, All0005, and Alr0523), inorganic ion transport and metabolism (All4121, Alr2877, and Alr3808), nucleotide transport and metabolism (all4131, Alr0051, and All2563), secondary metabolites biosynthesis, transport, and catabolism (All2655), and cell wall/membrane/envelope biogenesis (Alr4550, All4499, All3797, and Alr0834).

**Table 3.**
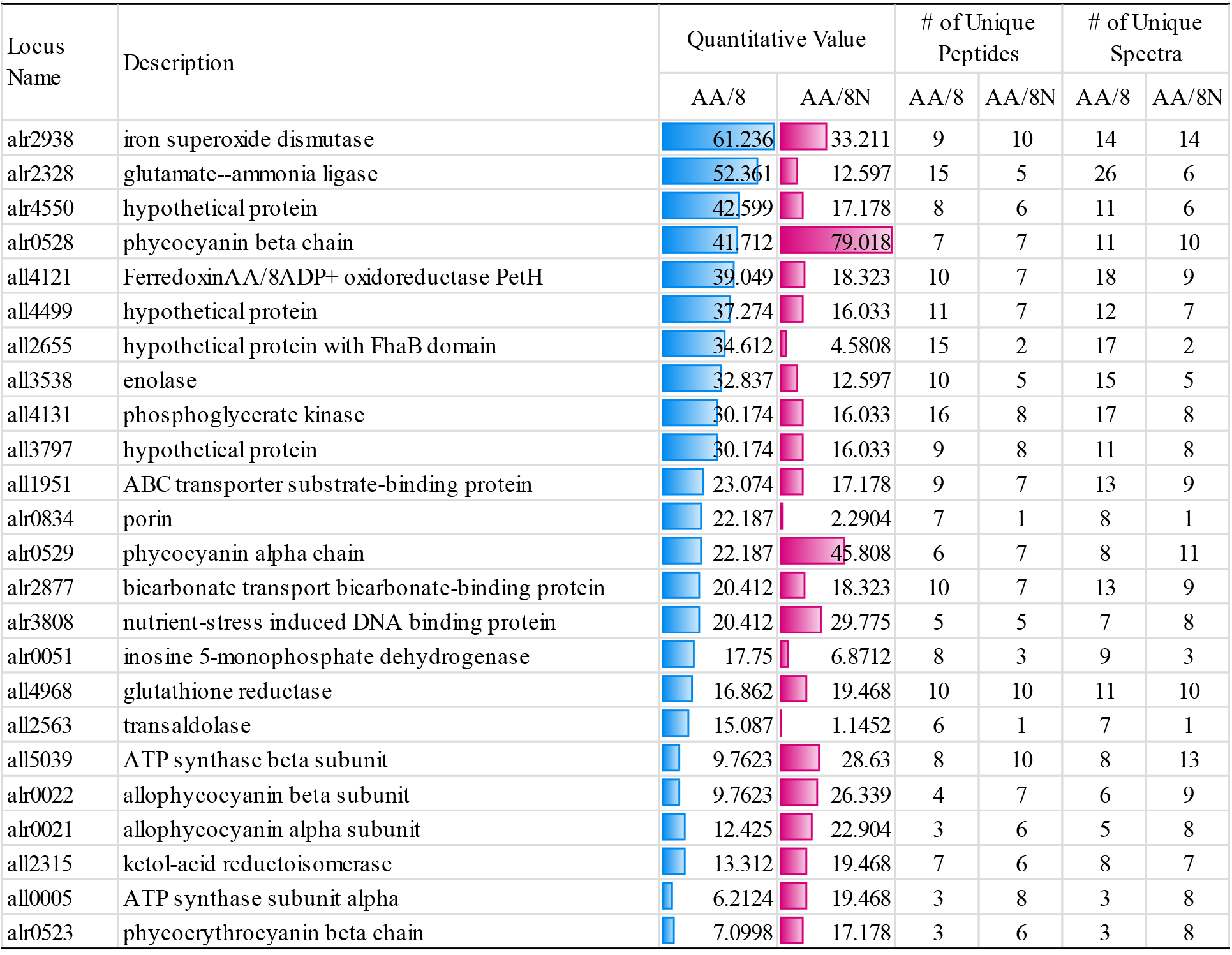
Top abundant proteins found in the culturing media with or without combined nitrogen (>15 quantitative Value)

Among the four exoproteins annotated to have function in cell wall/membrane/envelope biogenesis, Alr4550, All4499, and Alr0834 show homology with the outer membrane porin OprB of *Pseudomonas aeruginosa* (2, 31, 32). OprB is proposed to transport glucose into the periplasmic space, before further transported across the plasma membrane by an ABC-transporter (33). All4499, Alr4550 and Alr0834 are homology of 53% (Figure 4). Notably, the relative abundance of All4499 and Alr4550 in AA/8 *vs*. AA/8N culturing medium around 2.3 and 2.5-fold, respectively, Alr0834 showed 9.6-fold increase in *Anabaena sp*. PCC 7120 grown diazotrophically, agreeing with its up-regulation in responsible to nitrogen deprivation in both transcriptional and translational levels (24, 34). Taken together, these data suggest that Alr0834 plays a critical role in response to nitrogen stress under diazotrophically condition (35) and likely involved in selective transport of solutes and environmental resistance (10, 36), while All4499 and Alr4550 are notable for their high constitutive expression under standard lab oratory conditions. These OprB-like carbohydrate-selective porins act as channels in outer cell membrane, allowing carbohydrates and other molecules to pass into the cells (32).

**Figure 4.**
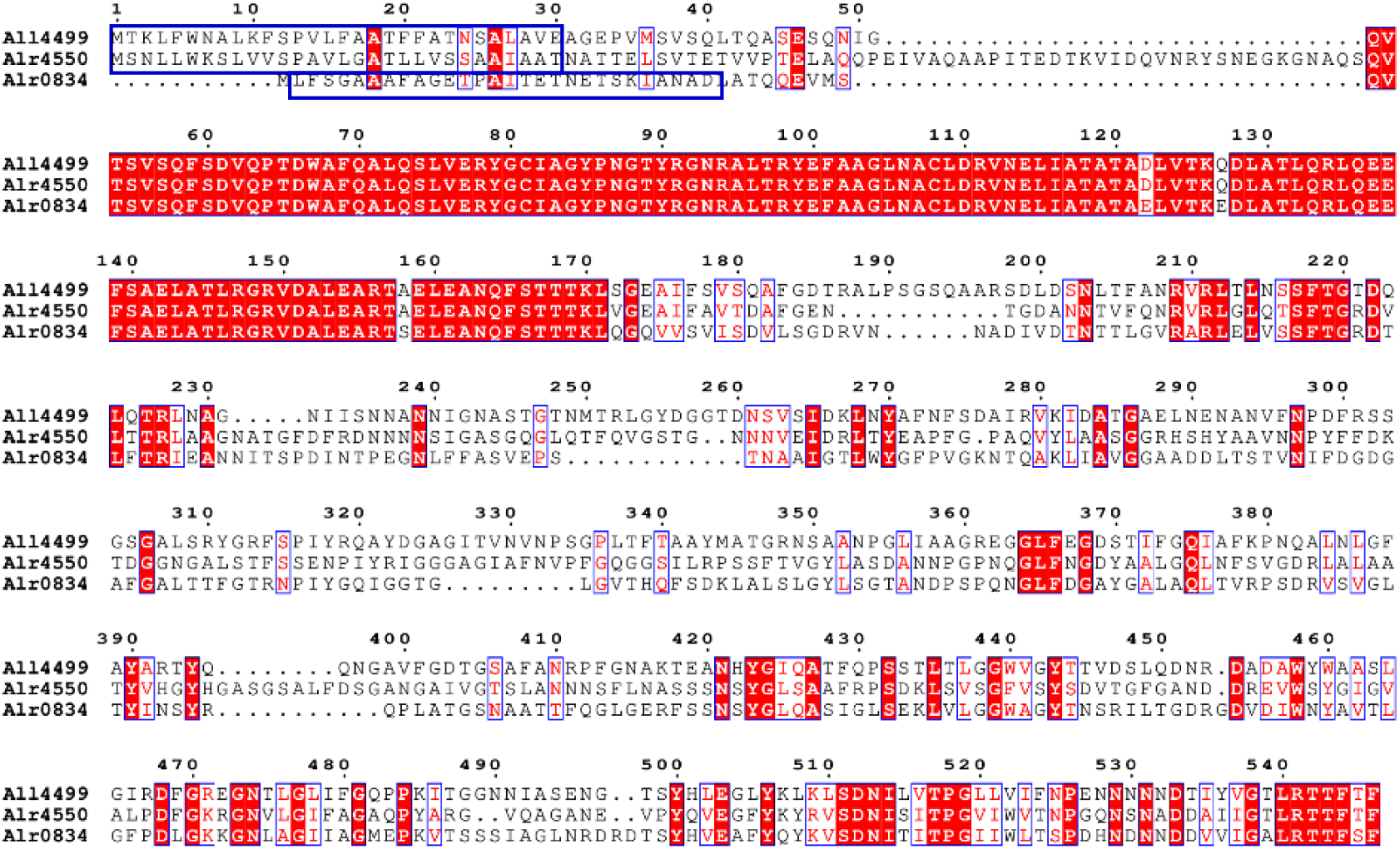
Multiple sequence alignment of All4499, Alr4550, and Alr0834 using ClustalW (https://www.genome.jp/tools-bin/clustalw) and the image was generated using ESPript 3.2 (https://espript.ibcp.fr/ESPript/ESPript/index.php). The N-terminal signal peptides were highlighted by blue boxes. All three proteins contain conserved S-layer homology domain.

### 3.5. Thirty-eight proteins predicted to be extracellular proteins using various computational prediction programs

To further understand the mechanism of these exoprotein secretion pathways, the putative signal peptide was analyzed using SignalP 6.0 program (https://services.healthtech.dtu.dk/services/SignalP-6.0/) (37). The putative signal peptides were identified in all three porin-related outer membrane proteins Alr4550, All4499, and Alr0834 (highlighted in Figure 4). Only 22% of all identified extracellular proteins contained a putative signal peptide in AA/8N and AA/8 culturing media (Table 4 and Table S3). There are 32 proteins identified and were annotated to be involved in cell wall/membrane/envelope biogenesis (All3797, All4388, All4499, Alr3345, Alr3539, Alr4550, Alr0834, and Alr2887), amino acid transport and metabolism (All1951, Alr0140, and Alr1834), carbohydrate transport and metabolism (All1027, All4539, and Alr4448), energy production and conversion (All0259 and Alr4251), inorganic ion transport and metabolism (All4575, All7121, Alr0608, and Alr2877), translation, ribosomal structure, and biogenesis (Alr1520), and posttranslational modification, protein turnover, chaperones (Alr0577).

**Table 4.**
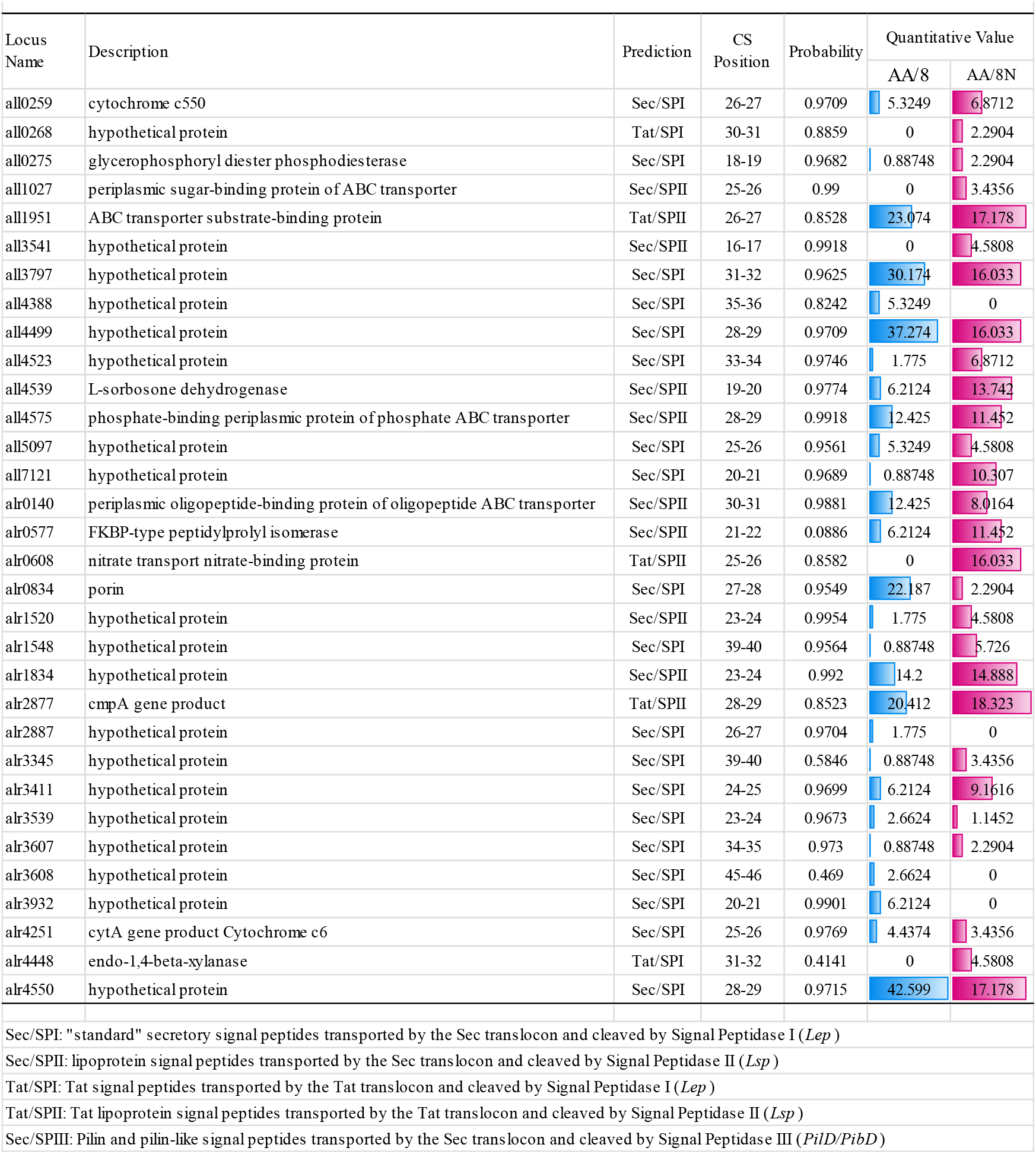
Prediction of the presence of signal peptides and the location of their cleavage sites in exoproteins identified by LC-MS/MS.

Among 32 exoproteins that predicted to have putative signal peptides, ten proteins were further predicted to be localized in extracellular or outer membrane or cell wall & surface (All0275, All7121, All4388, All4523, Alr3539, All4499, Alr0834, Alr2887, Alr4550, and Alr3345) using DeepLocPro 1.0 (https://services.healthtech.dtu.dk/services/DeepLocPro-1.0/) and/or pSORTb 3.0 (https://psort.org/psortb/). Six additional proteins were also predicted to be located either at cell wall & surface (All2655), or extracellular space (All4145, Alr0267, Alr4072, All1626, and Alr0996) using DeepLocPro 1.0 program (Table S3).

## 4. Conclusion

This study provides a comprehensive characterization of the extracellular proteome of *Anabaena sp*. PCC 7120 under nitrate-replete and diazotrophic growth conditions, revealing a pronounced nitrogen-dependent remodeling of extracellular protein composition. The observation that only a small proportion of extracellular proteins contain canonical signal peptides suggests that non-classical secretion pathways contribute substantially to protein export in this diazotrophic filamentous cyanobacterium.

Among the extracellular proteins identified, Alr0267 emerged as a particularly intriguing candidate. Its abundance under diazotrophic conditions, coupled with its heterocyst-specific localization and enrichment at the heterocyst periphery, provides the first evidence for extracellular protein secretion associated with heterocysts. Given that heterocysts possess specialized polysaccharide and glycolipid envelope layers that establish the microoxic environment required for nitrogen fixation, the apparent export of Alr0267 raises important questions regarding how proteins traverse these unique cellular barriers.

Collectively, our findings reveal evidence for a previously unrecognized heterocyst-associated protein secretion pathway and expand current understanding of protein trafficking in differentiated cyanobacterial cells. Future studies aimed at elucidating the molecular mechanisms underlying Alr0267 export may uncover novel secretion systems linked to heterocyst development and function. Beyond their biological significance, these discoveries may facilitate the development of *Anabaena* as a sustainable photosynthetic platform for the extracellular production of recombinant proteins and other value-added bioproducts powered by sunlight, atmospheric CO_2_, N_2_, and mineralized water.

## Supporting information

**S1**. Mascot Search Results for the specific protein bands in Figure 1

**S2**. Protein and Peptide Report for condense gel bands in Figure 2

**Figure S1**. Construction of pZR2099 plasmid containing P_*alr0267*_-Alr0267-GFP

**Table S1**. List of proteins identified in the exoproteome of *Anabaena* sp. PCC 7120 grown in nitrogen-fixing conditions (AA/8) or in media supplemented with nitrate (AA/8N).

**Table S2**. Comparison of the proteins identified in this work with previously reported exoproteomes of *Anabaena*. The exoproteins identified from *Anabaena sp*. PCC 7120 grown in AA/8 and AA/8N with the previously published exoproteomes of *Anabaena* sp. PCC 7120 from Sarasa-Buisan et al. 2024, Hahn et al., 2015 in BG11 medium, and Oliveira et al. 2015 in BG110, Bg11, and BG110+NH_4_+, respectively.

**Table S3**. Prediction of the exoproteins identified in *Anabaena sp*. PCC 7120 grown in nitrogen-fixing conditions (AA/8) or in media supplemented with nitrate (AA/8N).

## Availability of Data and Materials

All data generated or analyzed during this study are included in this published article. Further details or supporting materials are available from the corresponding author upon reasonable request.

## Ethics declarations

### Ethics Approval and Consent to Participate

Not applicable.

### Consent for publication

Not applicable.

## Acknowledgements

This research was supported by the National Science Foundation EPSCoR program, Collaborative Research: E-RISE RII: BioNitrogen Research Center (Award Number 2416911 to R. Z) and by NIFA grant “Genetic engineering of cyanobacteria to produce high-value proteins using atmospheric N_2_ gas” (Award number 2023-67022-39594) and partially by the South Dakota Agricultural Experiment Station-USDA Hatch project SD00H833-25.

## Author Contributions

Taufiq Nawaz contributed to data acquisition, data analysis, and drafted the first version of the manuscript. Ping He contributed to data acquisition and manuscript writing. Liping Gu contributed to experiment design, data analysis, LC–MS/MS data interpretation and led the manuscript revision. James Young contributed data analysis, bioinformatics, and computational prediction. Ruanbao Zhou conceived the project, provided overall supervision, and critically revised the manuscript. All authors read and approved of the final manuscript.

